# Estimating Protein Complex Model Accuracy Using Graph Transformers and Pairwise Similarity Graphs

**DOI:** 10.1101/2025.02.04.636562

**Authors:** Jian Liu, Pawan Neupane, Jianlin Cheng

## Abstract

**Motivation:** Estimation of protein complex structure accuracy is an essential step in protein complex structure prediction and is also important for users to select good structural models for various applications, such as protein function analysis and drug design. Despite the success of structure prediction methods such as AlphaFold2 and AlphaFold3, predicting the quality of predicted complex structures (structural models) and selecting top ones from large model pools remains challenging.

**Results:** We present GATE, a novel method that uses graph transformers on pairwise model similarity graphs to predict the quality (accuracy) of complex structural models. By integrating single-model and multi-model quality features, GATE captures both the characteristics of individual models and the geometric similarity between them to make robust predictions. On the dataset of the 15th Critical Assessment of Protein Structure Prediction (CASP15), GATE achieved the highest Pearson’s correlation (0.748) and the lowest ranking loss (0.1191) compared to existing methods. In the blind CASP16 experiment, GATE was ranked 4th according to the overall sum of z-scores of multiple metrics based on both TM-score and Oligo-GDTTS scores. In terms of per-target average metrics based on TM-score, GATE achieved a Pearson’s correlation of 0.7076 (1st place among all methods), a Spearman’s correlation of 0.4514 (3rd place), a ranking loss of 0.1221 (3rd place), and an Area Under the Curve (AUC) score of 0.6680 (3rd place), highlighting its strong, balanced ability of estimating complex model accuracy and selecting good models.

**Availability:** The source code of GATE is freely available at https://github.com/BioinfoMachineLearning/GATE.

## 1. Introduction

Proteins are fundamental to biological processes, and their three-dimensional (3D) structures determine their functions and interactions with other molecules. Accurate knowledge of protein structures is crucial for biomedical research and technology development. However, experimental structure determination methods like X-ray crystallography, nuclear magnetic resonance (NMR), and cryo-electron microscopy, while effective, are time-consuming and expensive. Consequently, computational protein structure prediction is essential for obtaining protein structures at a large scale.

Recent advances in deep learning have revolutionized protein structure prediction[1, 2, 3, 4, 5]. Methods such as AlphaFold2[1, 2] and AlphaFold3[3] can predict high-accuracy protein structures for most single-chain proteins (monomers) and a significant portion of multi-chain proteins (multimers / complexes). However, estimating the quality of the predicted structural models without ground-truth structures and identifying the best candidates from a pool of models (decoys) remains a significant challenge.

AlphaFold itself assigns an estimated quality score (e.g., pLDDT score or pTM score) to each structural model that it generates. However, it still cannot always assign higher estimated quality scores to better structural models[6]. Therefore, there is a significant need to develop independent protein model accuracy estimation (or quality assessment) methods to predict the quality of predicted protein structural models. These methods are not only important for ranking and selecting predicted protein structures but also crucial for determining how to use them appropriately in specific applications such as protein function analysis and protein structure-based drug design.

Traditionally, the methods for estimating protein model accuracy (EMA) can be classified in different ways. From the output of the methods, they can be classified as global quality assessment methods of assigning a single, global score to quantify the overall accuracy of a model and local quality assessment methods of assigning a quality score to each residue (or some specific residues such as interface residues in protein complexes). In this study, we focus on global quality assessment.

From the input, the EMA methods can be divided into single-model and multi-model methods. Single-model methods evaluate each model based on its own intrinsic properties, using statistical energy functions or machine learning, without comparing it with other models. Some examples of statistical energy-based methods include Rosetta[7], which rely on physical principles or statistical potentials. These methods are usually fast but cannot reliably correlates model energy and model quality due to the high complexity of protein energy landscape.

Machine learning-based single-model methods improve the accuracy of predicting model quality by integrating structural and sequence features of a model such as secondary structure and residue-residue contacts. Some early methods of using hand-crafted structural features to estimate the accuracy of protein tertiary (monomer) structures include ProQ[8], ModelEvaluator[9], ModFOLD[10], and QAcon[11].

More recently, deep learning-based methods, such as QDeep[12] and DISTEMA[13] were developed to learn features directly from raw structural data such as residue-residue distance maps or 3D atom grids of protein tertiary structures to estimate their quality. Particularly, some graph neural network (GNN)-based approaches, such as EnQA[6] and GCPNet-EMA[14], represent protein structures as graphs and integrate local and global structural information, outperforming traditional single-model approaches. Different from previous methods of predicting the quality of either tertiary structures or quaternary structures, both EnQA and GCPNet-EMA can be applied to predict the quality of both tertiary and quaternary structures, even though they were trained to predict the quality of tertiary structures only. The reason is that the graph representation they employ can be used to represent tertiary and quaternary structures well without any change.

Different from single-model EMA methods, multi-model EMA methods compare each model in a model pool with other models and rely its similarity with other models to estimate model quality. Therefore, they are often called consensus methods. One simple such approach is to use the average pairwise similarity between a model with all other models as its quality score, which is used by DeepRank3 for estimating quality of protein tertiary (monomer) structural models and by MULTICOM qa[15] for estimating the quality of protein quaternary (complex / multimer) structural models. Despite of the simplicity, on average, multi-model EMA methods still performed better than single-model EMA methods because in most situations many good models can be generated, leading to good consensus assessment[15, 16, 17]. However, multi-model EMA methods often fail for some hard targets when there are only a few good models in a large model pool or when there are many bad (or suboptimal) and similar models in the pool. Particularly, a large portion of bad, similar models always lead to bad consensus evaluation. In contrast, the performance of single-model methods is much more robust against the composition of models in a model pool because they provide an independent assessment of the quality of each individual model without considering the similarity between models. Therefore, single-model EMA methods are not susceptible to the influence of many bad and similar models and still have a chance to pick rare, good models.

To combine the strengths of single-model and multi-model EMA methods and address their limitations, we introduce GATE, a novel approach for predicting global protein structure quality using graph transformers on pairwise model similarity graphs. In this study, we focus GATE on predicting the global quality of protein quaternary structures instead of tertiary structures because there are much fewer EMA methods for quaternary structures and predicting quaternary structures is a much more challenging problem than the largely solved tertiary structure prediction problem. By capturing the relationships between models using model similarity graphs as well as the quality features of individual models represented as nodes in the graphs, GATE is able to make robust quality predictions.

## 2. Materials and Methods

The overall pipeline of GATE is illustrated in Figure 1. The approach integrates pairwise similarity graph construction, structural similarity-based subgraph sampling, node-level quality prediction using a graph transformer, and the aggregation of quality scores across subgraphs to predict the quality of each structural model (called decoy) in a model pool.

**Fig. 1.**
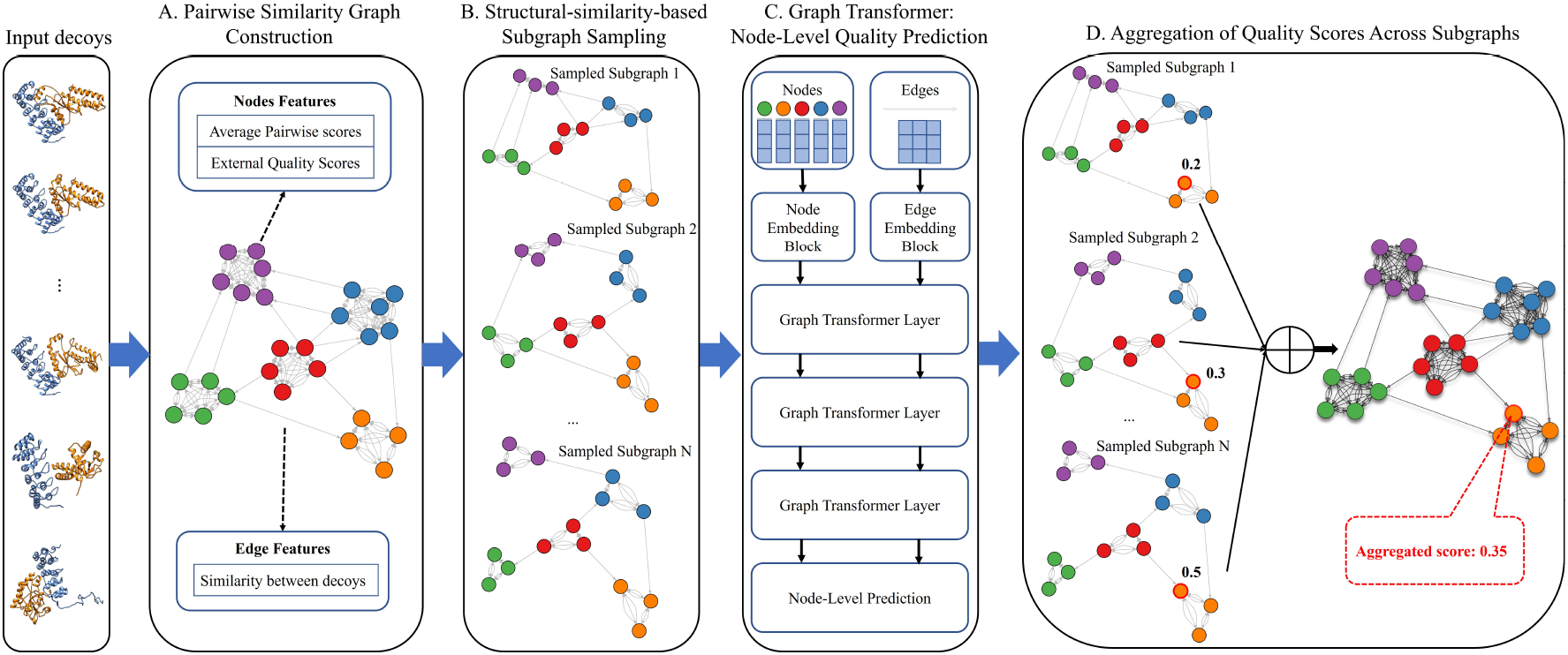
The workflow of GATE for predicting protein complex structure quality. The input consists of a set of protein complex structures (decoys) predicted from a protein sequence. (A) A pairwise similarity graph is constructed, where nodes represent individual decoys, and edges connect two structurally similar decoys. (B) Subgraphs are sampled based on structural similarity to make sure each group of similar models is equally represented in the subgraphs, preventing large groups from dominating small groups. (C) The sampled subgraphs are processed by a graph transformer to predict the quality score for each decoy. (D) Predicted quality scores from all sampled subgraphs are aggregated to produce the final quality score for each decoy.

### 2.1 Pairwise similarity graph construction

A pairwise similarity graph (Figure 1A) is constructed to captures the structural relationships between decoys. Each node represents an individual decoy, and an edge connects two decoys if they are similar according to a structural similarity metric such as TM-score, DockQ score[18], QS-score, and CAD-score[19]. To ensure meaningful structural relationships, in this study, an edge is established between two nodes if their TM-score exceeds 0.5. Even though only TM-score, a global similarity metric, is used to determine if there is an edge between two nodes, other complementary metrics such as QS-score of measuring inter-chain interface accuracy can also be used as the features of edges.

### 2.1 Structural similarity-based subgraph sampling

To prevent over-populated structurally similar low-quality models to dominate model quality assessment, which often happens with the consensus EMA methods, subgraphs are sampled from the full pairwise similarity graph to allow decoys of different structural folds are more equally represented. The sampling strategy is based on the structural similarity of the decoys in the model pool, quantified by the average pairwise similarity score (i.e., TM-score) (Figure 1B):

If the average pairwise similarity score is *<* 0.8, K-means clustering is applied to cluster the decoys into clusters, where each decoy is represented by a vector of similarity scores between it and the models in the pool. The number of clusters is determined by the silhouette score. A subgraph is generated by randomly sampling an equal number of decoys from each cluster. Many subgraphs are sampled so that each decoy may be included in multiple sub-graphs. If the average pairwise similarity score is *≥* 0.8, which usually indicates easy cases, subgraphs are randomly sampled from the entire graph without decoy clustering.

Each subgraph contains up to 50 nodes to balance computational efficiency and the need for structural variation.

### 2.3 Graph transformer

The subgraphs are processed using a graph transformer, which predicts the quality score for each decoy (Figure 1C). The transformer employs multi-head attention mechanisms to iteratively update node and edge embeddings, enabling it to capture both structural characteristics and contextual relationships.

#### 2.3.1 Node feature embedding

Node feature embeddings represent the structural characteristics of each decoy in a graph. The features of each node are designed to capture both local and global structural information. Key features include:

- **Average pairwise similarity scores** between a decoy represented by a node and other decoys: including TM-score and QS-score, calculated at both full graph and subgraph levels. These scores provide a comprehensive view of the structural similarity between each decoy and others, which resembe the traditional consensus scores.
- **Single-model quality scores** including single-model quality scores calculated by ICPS[15], EnQA, DProQA[20], and VoroIF-GNN[21] for a decoy, enriching the feature space with the quality assessment for individual models. These scores are normalized by multiplying the raw quality score by the ratio between the length of the decoy and that of the native structure. This normalization penalizes shorter decoys, ensuring that the scores reflect the completeness and accuracy of the decoy relative to the native structure.

The raw features above are passed through a multi-layer perceptron (MLP) with activation functions (e.g., LeakyReLU) to transform them into high-dimensional embeddings that better capture nonlinear relationships. These embeddings serve as the initial representations for each decoy and are iteratively updated in the subsequent layers of the graph transformer.

#### 2.3.2 Edge feature embedding

Edge features encode pairwise relationships between decoys, initialized using the number of common interaction interfaces and similarity scores between two connected decoys, including TM-score and QS-score. These metrics capture global and interfacial similarity between connected nodes, respectively.

Like the node feature embedding, the raw edge features are mapped into a higher-dimensional space through an MLP. Each edge embedding is refined by the graph transformer through iterative updates, integrating information from the neighboring nodes. This process allows the edge embeddings to dynamically leverage the contextual relationships between decoys.

#### 2.3.3 Graph transformer layer

The graph transformer layers (Figure 1C) iteratively update node and edge embeddings through multi-head attention and feed-forward neural networks. The attention mechanism uses the node embeddings to generate queries (Q), keys (K), and values (V), while the edge embeddings are incorporated as biases to modulate pairwise interactions between nodes. The attention scores are computed as:

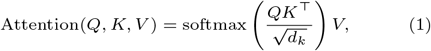

where *Q, K*, and *V* are the query, key, and value matrices, respectively, and *d*_*k*_ is the dimension of the key. To update the node embeddings, the model aggregates information from neighboring nodes using attention scores. For a node *i*, the updated embedding is calculated by:

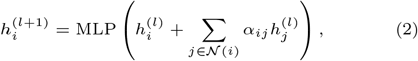

where 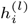 is the embedding of node *i* at layer *l, N* (*i*) represents its neighbors, and *α*_*ij*_ is the attention score between nodes *i* and *j*. The MLP introduces non-linearity, enabling the model to capture complex structural relationships.

The edge embeddings are updated similarly by integrating information from their associated nodes. The edge embedding between nodes *i* and *j* at layer *l* is updated as:

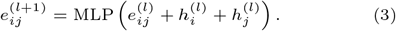

Residual connections and layer normalization are applied to both node and edge updates, ensuring stable training and efficient learning across layers.

By iteratively applying these updates, the graph transformer learns to capture intricate structural dependencies and relationships within the pairwise similarity graph, enabling it to effectively predict quality scores for the decoys.

#### 2.3.4 Node-level prediction

The graph transformer predicts a global quality score for each decoy (node) in an input subgraph. The target global quality score is the true TM-score between a decoy and the native structure, calculated by USalign[22].

#### 2.3.5 Loss Function

The graph transformer is trained using a combination of a Mean Squared Error (MSE) loss and a pairwise loss:

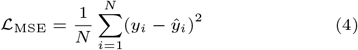

where *y*_*i*_ is the true quality score for decoy 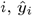 the predicted score, and N the number of decoys.

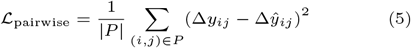

where *P* represents the number of decoy pairs,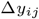 is the true difference between quality scores of decoys *i* and *j*, and 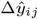 is the predicted difference. This loss enables the model to learn the relative quality between two models such that a better model has a higher score than a worse one.

The total loss is a weighted sum of the two losses above:

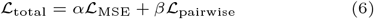

where *α* and *β* control the importance of each component.

### 2.4 Aggregation of predicted quality scores across subgraphs

The final predicted quality score for each decoy is aggregated across subgraphs using either mean or median aggregation:

- **Mean aggregation**: averaging predictions from all subgraphs containing the decoy.
- **Median aggregation**: using the median of predictions, which is robust to outliers.

### 2.5 Training, Validation and Test Procedure

The graph transformer was first trained, validated, and tested on the CASP15 dataset and then blindly tested in the CASP16 experiment. The CASP15 dataset comprises 41 complex targets, with each target having an average of 270 decoys. The CASP15 dataset was divided into training, validation, and test sets by targets using a 10-fold cross-validation strategy to enable the robust evaluation of the transformer’s performance. Specifically, it was split into 10 subsets, each containing 4–5 targets. For each target, a fixed number of 2,000 subgraphs were sampled from the full model similarity graph of all the decoys of the target, with each subgraph containing exactly 50 nodes. Each subset contains 8000–10000 subgraphs. In each round of training, validation, and testing, 8 subsets were used for training (parameter optimization), 1 for validation (model selection), and 1 for testing. The test results over the 10 subsets were pooled together and evaluated as a whole.

The training was performed using a Stochastic Gradient Descent (SGD) optimizer, with the hyperparameters organized into the following groups:

- **SGD optimizer parameters**, including the learning rate, weight decay, and batch size, which were tuned to ensure effective optimization and convergence during training.
- **Graph transformer parameters**, including the number of attention heads, the dimensionality of node and edge embeddings, the number of graph transformer layers, and the dropout rate. These parameters define the architecture and control regularization of GATE.

The trained GATE models were selected according to their performance on the validation set in terms of the combined loss function considering both the Mean Squared Error (MSE) loss and the pairwise loss.

Three different kinds of GATE models below were trained, validated, and tested on the same 10-fold data split:

- **GATE-Basic:** use the basic set of node and edge features described above.
- **GATE-GCP**: add the single-model quality score calculated by GCPNet-EMA for each deocy as an extra node feature.
- **GATE-Advanced**: Further augment the node and edge feature sets using advanced interface quality scores, such as DockQ ave[23], DockQ wave[23], and CAD-scores. Specifically, the average interface scores between a decoy and other decoys are used as its extra node features and the interface scores between the two nodes of an edge are used as its extra edge features.

## 3. Results

### 3.1 Evaluation metrics

The following metrics were used to assess the performance of the EMA methods:

- **Pearson’s correlation (Corr**^**p**^): Pearson’s correlation measures the linear relationship between the predicted and true quality scores. A higher Pearson’s correlation indicates a stronger alignment between the predicted and true values, reflecting a method’s accuracy in predicting overall structural quality.
- **Spearman’s correlation (Corr**^**s**^): Spearman correlation assesses the rank-order relationship between the predicted and true quality scores. This metric evaluates how well the predicted scores preserve the correct ranking of decoys, independent of the exact score values.
- **Ranking loss:** Ranking loss quantifies the ability of a method to correctly rank the best decoy in the pool. It is calculated as the difference between the true quality (e.g., TM-score) of the best decoy in the pool and the true quality of the decoy with the highest predicted quality. Lower ranking loss indicates better performance in identifying the top-quality structure.
- **Receiver Operating Characteristic (ROC) Curve and Area Under the Curve (AUC):** The ROC curve illustrates the trade-off between the true positive rate (sensitivity) and the false positive rate at various classification thresholds. By setting the threshold to the 75% quantile of true TM-scores of the decoys to divide them into two classes: positive ones and negative ones, we make the metric focus on identifying better-quality decoys. The AUC provides a single scalar value summarizing the ROC curve, with higher values indicating better discrimination. An AUC of 1.0 represents perfect classification, while an AUC of 0.5 indicates random performance. This metric is particularly useful for assessing a method’s robustness in ranking decoys and identifying top-quality structures.

Each metric above is calculated for the decoys of each target and then average over all the targets in a dataset.

### 3.2 Performance of GATE models on the CASP15 dataset

The performance of three GATE models and their ensemble (GATE-Ensemble) in comparison with several existing methods on the CASP15 dataset is reported in Table 1. Among the individual models, GATE-Basic, using the basic features, achieved a Pearson’s correlation (Corr^p^) of 0.7447, a ranking loss of 0.1128, and an AUC of 0.7181. GATE-GCP, which included an additional GCPNet-EMA feature, showed a slight improvement in Spearman correlation (0.5788) over GATE-Basic (0.5722) but had similar Pearson’s correlation (0.7454) and slightly worse ranking loss (0.1186), indicating that the added feature had minimal impact on the performance.

**Table 1.** Comparison of GATE models and other methods in terms of Pearson’s correlation (Corr^p^), Spearman’s correlation (Corr^s^), ranking loss, and AUC on the CASP15 complex structure dataset. The term norm indicates that the quality scores predicted by a method are normalized by the length of the predicted structure relative to the native structure. Only the performance of the normalization version of such a method is shown because it works better than the unnormalized version. Bold font denotes the best result, while the second best result is underlined.

| Method | $\text{Corr}^P$ | $\text{Corr}^S$ | Ranking loss | AUC |
| --- | --- | --- | --- | --- |
| PSS | 0.7292 | <u>0.5755</u> | 0.1406 | 0.7137 |
| AlphaFold pLDDT<sub>norm</sub> | 0.2578 | 0.2611 | 0.1793 | 0.6399 |
| DProQA<sub>norm</sub> | 0.1598 | 0.1174 | 0.2555 | 0.5942 |
| EnQA<sub>norm</sub> | 0.2898 | 0.2139 | 0.2188 | 0.5965 |
| VoroIF-GNN-score<sub>norm</sub> | 0.1972 | 0.0966 | 0.2092 | 0.5694 |
| Average-VoroIF-GNN-residue-pCAD-score<sub>norm</sub> | 0.1335 | -0.0027 | 0.1737 | 0.5526 |
| VoroMQA-dark global score<sub>norm</sub> | 0.0253 | 0.0037 | 0.1265 | 0.5580 |
| GCPNet-EMA<sub>norm</sub> | 0.3216 | 0.2696 | 0.2052 | 0.6379 |
| GATE-Basic | 0.7447 | 0.5722 | <u>0.1128</u> | 0.7181 |
| GATE-GCP | <u>0.7454</u> | **0.5788** | 0.1186 | <u>0.7191</u> |
| GATE-Advanced | 0.7224 | 0.5416 | **0.1018** | 0.6981 |
| GATE-Ensemble | **0.7480** | 0.5754 | 0.1191 | **0.7194** |

GATE-Advanced, with addition of several interfacial quality features like DockQ ave and DockQ wave, achieved the lowest ranking loss (0.1018), excelling in identifying top-quality decoys. However, its Pearson’s correlation (0.7224) was slightly lower than that of GATE-Basic and GATE-GCP, suggesting a tradeoff between the correlation and the ranking loss. This result highlights that while GATE-Advanced can select the top-1 model better (minimizing the ranking loss), its absolute quality predictions may be less precise.

GATE-Ensemble, which averages predictions from GATE-Basic, GATE-GCP, and GATE-Advanced, outperformed the three component methods in terms of Pearson’s correlation and AUC score but underperformed them in terms of the ranking loss. Specifically, it achieved the highest Pearson’s correlation (0.7480), a balanced Spearman correlation (0.5754), a ranking loss of 0.1191, and the highest AUC of 0.7194. Overall, ensembling the three component methods gained marginal improvement.

All the GATE models substantially outperformed single-model methods, including AlphaFold plDDT_norm_, DProQA_norm_, EnQA_norm_, VoroIF-GNN-score_norm_, Average-VoroIF-GNN-residue-pCAD-score_norm_, VoroMQA-dark global score_norm_, and GCPNet-EMA_norm_ in terms of all the metrics in almost all the cases. The results demonstrate that integrating single-model quality scores and similarity between decoys in GATE substantially improves the estimation of the accuracy of decoys.

The consensus method based on the average pairwise similarity score (PSS) between decoys calculated by USalign also performed generally better than the single-model EMA methods. Its performance in terms of Pearson’s correlation, Spearman’s correlation, and AUC is comparable to the GATE models, but its ranking loss is substantially higher (worse) than them. For instance, the ranking loss of PSS is 0.1406, which is much higher than 0.1018 of GATE-Advanced, 0.1128 of GATE-Basic, 0.1186 of GATE-GCP, and 0.1191 of GATE-Ensemble. A direct comparison of GATE-Ensemble and PSS shows that GATE-Ensemble outperformed PSS in terms of Pearson’s correlation, ranking loss and AUC, while they performed almost the same in terms of Spearman’s correlation. The biggest difference between the two lies in Pearson’s correlation and ranking loss. The per-target Pearson’s correlation and ranking loss comparison between the two methods is visualized in supplementary Figure S1A and S1B, respectively. In terms of ranking loss, GATE-Ensemble performed much better on five targets (H1114, H1144, H1167, T1115o, and T1161o) that are highlighted in different colors in supplementary Figure S1 and much worse than PSS on only one target (T1181o).

However, a detailed analysis of T1181o (a homo-trimer) revealed that the top-1 decoys selected by PSS and GATE-Ensemble are very similar (TM-score between them = 0.89), suggesting that their true quality scores should also be comparable. However, when aligning each decoy with the true structure, USalign successfully identifies an optimal chain mapping for the decoy selected by PSS, assigning it a high TM-score. In contrast, it fails to find an optimal chain mapping for the decoy selected by GATE-Ensemble, resulting in a low TM-score and, consequently, a high loss. This discrepancy indicates that the high loss observed for GATE-Ensemble on T1181o is not due to model selection but rather stems from a chain mapping issue within USalign. Although this issue only arises occasionally for certain targets, it necessitates manual inspection until the chain mapping algorithm in USalign is improved[4].

The histogram of the true TM-scores of the decoys of the five targets on which GATE-Ensemble substantially outperformed PSS are plotted in supplementary Figure S1C. One common pattern among the five targets is that common decoys of high frequency in the histograms have mediocre scores.

For a target H1114 ([NiFe]-hydrogenase, oxidoreductase, energy metabolism, stoichiometry: A4B8C8; skewness of the distribution of the TM-scores of the decoys = 0.831), GATE-Ensemble achieved a ranking loss of 0.0398 and a Pearson’s correlation of 0.722, outperforming PSS (ranking loss 0.2520, correlation 0.393). Similarly, for target H1144 (nanobody, A1B1; skewness = -1.176), GATE-Ensemble excelled with a ranking loss of 0.0017 and a correlation of 0.755, compared to PSS (ranking loss 0.3117, correlation 0.485).

For target H1167 (antibody-antigen complex, stoichiometry: A1B1C1; skewness = 0.024), representing a balanced decoy pool, GATE-Ensemble achieved a ranking loss of 0.0318 and a correlation of 0.702, far surpassing PSS (ranking loss 0.3095, correlation 0.335). Target T1115o (A16, skewness = 0.567) followed a similar trend, with GATE-Ensemble (ranking loss 0.1156, correlation 0.702) outperforming PSS (ranking loss 0.3668, correlation 0.465).

For the highly skewed target T1161o (Dimeric DZBB fold, DNA-binding protein, stoichiometry: A2; skewness = 2.258), GATE-Ensemble achieved a perfect ranking loss of 0 and a correlation of 0.740, compared to PSS (ranking loss 0.4692, correlation 0.362), demonstrating its robustness in selecting good decoys from in decoy pools dominated by low-quality decoys.

In summary, GATE-Ensemble consistently outperformed PSS in both ranking loss and Pearson’s correlation across all five targets because PSS selected common decoys of low or mediocre quality (i.e., the central tendency problem), while GATE-Ensemble is able to overcome the problem by using its graph transformer architecture and sub-graph sampling to consider both the quality of individual decoys and the similarity between them.

### 3.3 Performance of GATE in the blind CASP16 experiment

During CASP16, GATE blindly participated in the EMA category under the predictor name MULTICOM GATE. The 10 models for each of three GATE variants (GATE-Basic, GATE-GCP, and GATE-Advanced) trained on the CASP15 dataset via the 10-fold cross-validation, i.e., 30 models in total, were ensembled together to make prediction. MULTICOM GATE used the average output of the 30 models as prediction. Moreover, to ensure stable predictions given the randomness in subgraph sampling, MULTICOM GATE made predictions for the decoys of each target five times and averaged them as the final prediction.

Out of the 38 CASP16 complex targets evaluated, GATE models were used to make predictions for 36 targets, with H1217 and H1227 excluded due to the limitation of computational resource. The performance of MULTICOM GATE in comparison with other CASP16 EMA methods on the 36 targets is illustrated in supplementary Figure S2 and summarized in Table 2, using two perspectives: the sum of z-scores (supplementary Figure S2) and per-target-average metrics (Table 2).

**Table 2.** Average per-target evaluation metrics (Pearson’s correlation, Spearman’s correlation, ranking loss and AUC) of 23 CASP16 predictors in terms of TM-score. The best performance for each metric is shown in bold, the second-best is underlined, and the third-best is underlined and italicized. The methods are ordered by the ranking loss from low to high. MULTICOM GATE ranked among top three according to every metric.

| Predictor Name | Corr<sup>P</sup> | Corr<sup>S</sup> | Ranking loss | AUC |
| --- | --- | --- | --- | --- |
| GuijunLab-QA | 0.6479 | 0.4149 | **0.1195** | 0.6328 |
| MULTICOM | 0.6156 | 0.4380 | <u>0.1207</u> | 0.6660 |
| **MULTICOM_GATE** | **0.7076** | <u>0.4514</u> | <u>0.1221</u> | <u>0.6680</u> |
| MULTICOM_LLM | <u>0.6836</u> | **0.4808** | 0.1230 | <u>0.6685</u> |
| MIEnsembles-Server | 0.6072 | 0.4498 | 0.1325 | 0.6670 |
| GuijunLab-Pathreader | 0.5309 | 0.3744 | 0.1331 | 0.6237 |
| ModFOLDdock2 | <u>0.6542</u> | <u>0.4640</u> | 0.1371 | **0.6859** |
| ModFOLDdock2R | <u>0.5724</u> | 0.3867 | 0.1375 | 0.6518 |
| VifChartreuse | 0.2921 | 0.2777 | 0.1440 | 0.6149 |
| MQA_base | 0.4331 | 0.2897 | 0.1462 | 0.6085 |
| MQA_server | 0.4326 | 0.2913 | 0.1468 | 0.6120 |
| GuijunLab-Human | 0.6327 | 0.4148 | 0.1477 | 0.6368 |
| MULTICOM_human | 0.5897 | 0.4260 | 0.1518 | 0.6576 |
| ChaePred | 0.4548 | 0.3971 | 0.1580 | 0.6534 |
| AF_unmasked | 0.4015 | 0.2731 | 0.1595 | 0.6052 |
| VifChartreuseJaune | 0.3421 | 0.1756 | 0.1630 | 0.5951 |
| GuijunLab-Assembly | 0.5439 | 0.3280 | 0.1636 | 0.6191 |
| Guijunlab-Complex | 0.4889 | 0.3019 | 0.1792 | 0.6054 |
| ModFOLDdock2S | 0.5285 | 0.3116 | 0.1806 | 0.6084 |
| MULTICOM_AI | 0.3281 | 0.2623 | 0.1913 | 0.6057 |
| COAST | 0.3840 | 0.2297 | 0.2091 | 0.6072 |
| MQA | 0.4410 | 0.2425 | 0.2183 | 0.5858 |
| PIEFold_human | 0.1929 | 0.1451 | 0.2306 | 0.5497 |

Supplementary Figure S2 presents the weighted sum of the z-scores for each evaluation metric, including Pearson’s correlation (*P* (*TM* -*score, p*) in terms of TM-score, *P* (*Oligo*-*GDTTS, p*)) in terms of oligomer global distance test (GDT-TS) score, Spearman’s correlation (*S*(*TM* -*score, p*) in terms of TM-score, *S*(*Oligo*-*GDTTS, p*)) in terms of oligomer GDT-TS score, ranking loss (*L*(*TM* -*score, p*) in terms of TM-score, *L*(*Oligo*-*GDTTS, p*)) in terms of oligomer GDT-TS score, AUC (*R*(*TM* -*score, p*) in terms of TM-score, and AUC (*R*(*Oligo*-*GDTTS, p*)) in terms of oligomer GDT-TS score. The z-scores of a predictor for each target were calculated relative to the mean and standard deviation of all the CASP16 predictors. The per-target z-score are then summed across all 36 targets to quantify the overall performance of each predictor. This approach used by the CASP16 official EMA assessment provides a comprehensive evaluation of each predictor’s relative performance in terms of multiple complementary metrics.

MULTICOM GATE achieved the highest sum of z-scores for *P* (*TM* -*score, p*) and *P* (*Oligo*-*GDTTS, p*), demonstrating its consistent ability to predict structural quality across all targets. It also ranked among the top methods in *S*(*TM* -*score, p*) and *S*(*Oligo*-*GDTTS, p*), highlighting its robustness in ranking decoys accurately. Furthermore, MULTICOM GATE performed competitively in ranking loss (*L*(*TM* -*score, p*) and *L*(*Oligo*-*GDTTS, p*)), reflecting its effectiveness in identifying the highest-quality decoys. Its strong performance in *R*(*TM* -*score, p*) and *R*(*Oligo*-*GDTTS, p*) underscores its ability to distinguish between high- and low-quality decoys. According to the weighted sum of all the z-scores, MULTICOM GATE ranked no. 4 among the 23 CASP16 predictors.

It is worth noting that z-score amplifies the impact of a target on which a predictor performs very well while most other predictors perform poorly. Another standard way to evaluate the predictors is to simply compare their average scores across all the test targets, without emphasizing specific ones. Table 2 complements supplementary Figure S2 by reporting the original per-target average performance metrics in terms TM-score, which was what MULTICOM GATE was trained to predict. These metrics include Pearson’s correlation (Corr^p^), Spearman’s correlation (Corr^s^), ranking loss, and AUC, averaged across the 36 targets. MULTICOM GATE achieved the highest Corr^p^ (0.7076), demonstrating its strong capability to predict structural quality based on TM-score. It also ranked third in Corr^s^ (0.4514) and third in ranking loss (0.1221), reflecting its balanced performance in both ranking consistency and identifying top-quality decoys. Additionally, MULTICOM GATE achieved a competitive AUC value (0.6680), ranking third. Overall, MULTICOM GATE delivered a balanced, excellent performance, ranking among top three in terms of every metric.

To illustrate how GATE identifies high-quality decoys, supplementary Figure S3 presents the pairwise similarity graph for a hard CASP16 target H1215, which is a heterodimer consisting of a mNeonGreen protein (antigen) bound with a nanobody. The color of nodes in the graph corresponds to the true TM-scores of the decoys represented by them, ranging from 0.5 (yellow) to 1.0 (purple).

A low-quality but common decoy, H1215TS196 4 (true TM-score = 0.613, ranking loss = 0.377), was selected as the top-1 decoy by the average pairwise structural similarity (consensus) strategy. The decoy has an incorrect interface between its two subunits and has a high loss. In contrast, a high-quality decoy with a correct interface, (H1215TS014 2 (true TM-score = 0.987, ranking loss = 0.003), was identified by MULTICOM GATE as top 1. The success of MULTICOM GATE on this target is significant as many CASP16 EMA predictors failed on this target. This example demonstrates that GATE effectively integrates local and global structural features to prioritize high-quality decoys over common mediocre models selected by the consensus approach. By leveraging both pairwise similarity information and single-model quality scores, GATE accurately distinguishes the decoys with near-native fold, as indicated by the high TM-score of the selected decoy.

Together, the results above provide a holistic view of MULTICOM GATE’s performance. The sum of z-scores reflect its relative strength across the targets, while the per-target-average metrics validate its absolute performance for TM-score prediction. These results demonstrate that MULTICOM GATE effectively integrates local and global structural features to deliver robust and accurate predictions, solidifying its position as one of the top-performing methods in the blind CASP16 experiment.

### 3.4 Limitation

While GATE demonstrates strong performance in our own test and the blind CASP16 experiment, it still has some limitations. First, generating pairwise similarity scores for a large number of decoys of a large protein complex is a time-intensive process because the complex structure comparison tools used in this study, such as MMalign and USalign, are slow in comparing large mutli-chain protein structures. A potential solution is to use faster approximate methods, such as FoldSeek[24], to accelerate structural comparisons.

Second, subgraph sampling introduces some randomness so that running GATE multiple times may get slightly different quality scores for a decoy. However, this effect can be minimized by extensive sampling of sub-graphs and averaging the predictions. In our experiment, if 2,000 subgraphs are sampled and their predictions are aggregated to produce a final score, the aggregation smooths out variability, ensuring robust and consistent quality scores are produced even for diverse or imbalanced decoy pools. Consequently, the randomness from sampling has a negligible impact on the overall performance of GATE. Moreover, the randomness also has a positive effect because it allows GATE to generate standard deviations for predicted scores.

## 4. Conclusion

This study introduces a novel graph-based approach to improve protein complex structure quality assessment by leveraging pairwise similarity graphs and graph transformers to integrate multiple complimentary features and the strengths of multi-model and single-model EMA methods. The approach was rigorously tested on the CASP15 dataset as well as in the blind CASP16 experiment, demonstrating the robust and promising performance across multiple evaluation metrics. Particularly, it maintains the advantage of the multi-model consensus methods over the single-model methods, while overcoming the failure of the central tendency of the consensus approach in some cases. In the future, we plan to speed up the construction of pairwise similarity graphs and explore more advanced graph neural network architectures and larger training datasets to better integrate structural and similarity features to further enhance the performance of the approach.

## Author contributions statement

J.L. contributed to writing, software development, visualization, formal analysis, data curation, and validation. P.N. contributed to formal analysis. J.C. was responsible for conceptualization, method design, investigation, funding acquisition, writing, validation, software, formal analysis, project administration, supervision, resources, and visualization.

## Acknowledgments

This work is supported in part by a NSF Grant (Award Number: DBI2308699) and a NIH grant (Award Number: R01GM093123).

