## Supplementary material for "Estimating Protein Complex Model Accuracy Using Graph Transformers and Pairwise Similarity Graphs": Supplmentary material

February 7, 2025

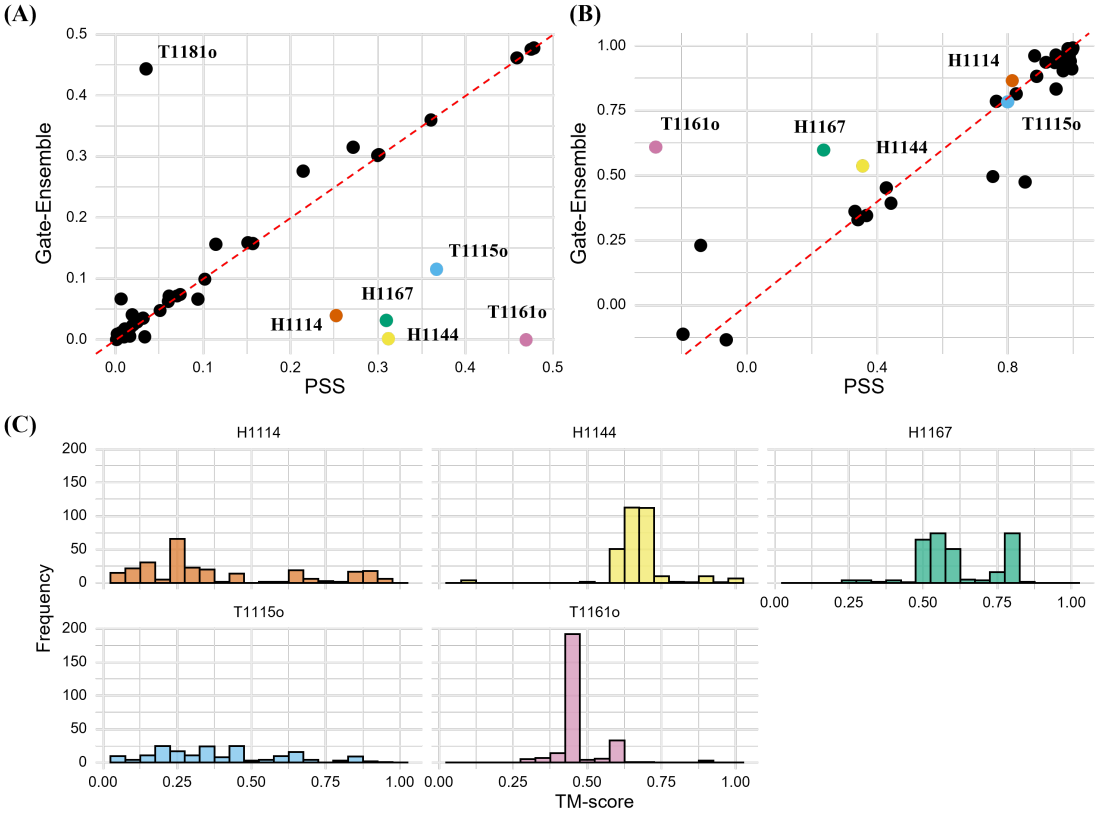


Figure S1: Per-target performance comparison between the PSS and GATE-Ensemble on CASP15 complex targets. (A) Ranking loss of GATE-Ensemble plotted against that of PSS; (B) Pearson's correlation of GATE-Ensemble plotted against that of PSS; (C) the histogram (distribution) of true TM-scores of the decoys for H1114, H1144, H1167, T1115o, T1161o.

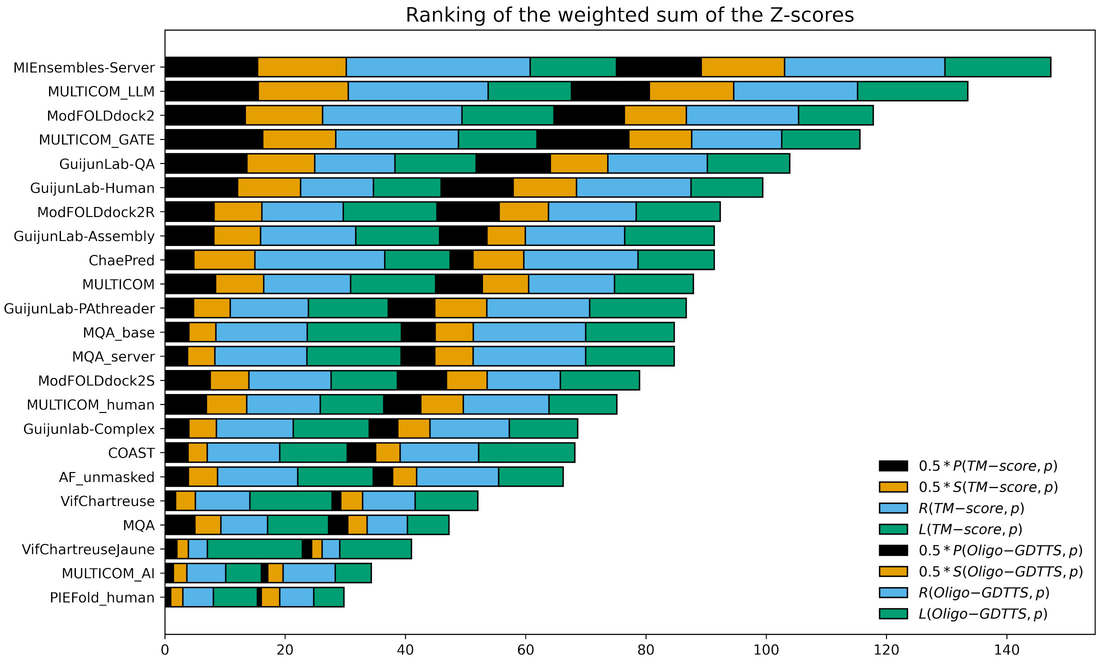


Figure S2: The overall performance of 23 CASP16 predictors in estimating the accuracy of the structural models of 36 CASP16 multimer targets according to the z-scores of multiple evaluation metrics (i.e., Pearson's correlation, Spearman's correlation, AUC, and ranking loss) in terms of both TM-score and oligomer GDT-TS score. Each kind of z-score is denoted by a colored bar. The predictors are ordered according to the weighted sum of all the z-scores.

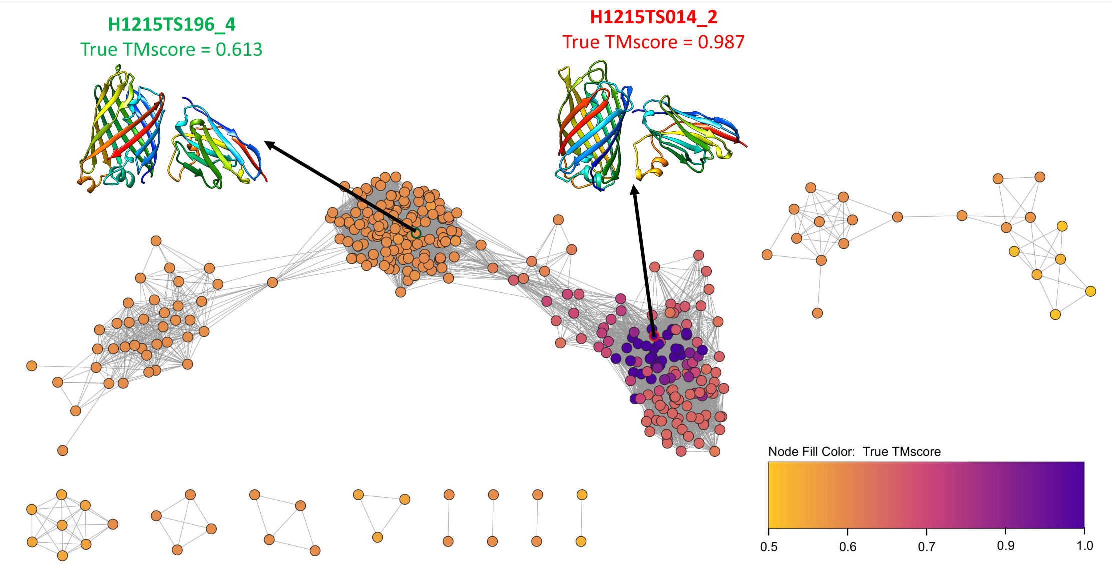


Figure S3: The pairwise similarity graph for the structural modes (decoys) of CASP16 target H1215. The top 1 decoy selected by the average pairwise similarity score (H1215TS196_4, true TM-score = 0.613) and MULTICOM_GATE (H1215TS014_2, true TM-score = 0.987).
